# Generation and characterisation of an estrogen receptor-positive GEMM-derived *Pten p53* null transplantable breast tumour model for therapeutic testing

**DOI:** 10.1101/762047

**Authors:** E.J. Davies, H. Morgan, G. Tornillo, C.D. Chabbert, H. Kendrick, M.J. Adhesmaki, S. Luke, S.T. Barry, M.J. Smalley

**Author notes:** Intertek Pharmaceutical Services Manchester, Hexagon Tower, Blackley, Manchester, M9 8GQ (H. Kendrick). Roche Innovation Center Zürich, Wagistrasse 10, 8952 Schlieren, Switzerland (C.D. Chabbert). Authors made equal contributions (E.J. Davies, H. Morgan). Authors made equal contributions (S.T. Barry, M.J. Smalley).

## Abstract

Assessing the signalling pathway dependencies of tumours that arise autochthonously in genetically engineered mouse models (GEMMs) of breast cancer is particularly challenging due to the high degree of intra- and inter-tumour heterogeneity, as well as the long latency of tumour development in such models. Use of transplantable tumour lines derived from autochthonous tumours (‘Mouse Derived Xenografts’ or MDXs) is one possible solution and has been used successfully in models of BRCA1-associated triple negative breast cancer. However, their potential in ER+ breast cancer models has not been addressed. Here, we assess the utility of orthotopic transplantable tumour lines derived from an autochthonous ER+ *Blg-cre Pten^fl/fl^ p53^fl/fl^* breast cancer model. We show that initial tumour implantation and early passage results in the development of lines of progeny with heterogeneous histopathological phenotypes which is coincident with an accumulation of, or selection for, *de novo* mutations. Importantly, these lines also display different dependencies on the key pathways that drive tumourigenesis, which can lead to inherent resistance to treatment with pharmacological agents targeting these pathways and makes them important models to test strategies to overcome such resistance.

## Introduction

Increasing complexity of genetically engineered mouse tumour models (GEMMs) and their improved representation of human pathology and pharmacokinetics means that they are increasingly important in understanding tumour biology and response to therapeutics (Gopinathan et al., 2015, van Miltenburg and Jonkers, 2012, Combest et al., 2012, Usary et al., 2016). In breast cancer, GEMMs have been useful tools in deconvoluting the specific contribution of mutation and cell of origin in tumour development (Melchor et al., 2014, Molyneux et al., 2010, Molyneux and Smalley, 2011). However, the use of these advanced autochthonous models for therapeutic testing is challenging as the tumours that develop in a single model are often heterogeneous, recapitulating a range of breast cancer subtypes observed in humans, and often at long latencies. One approach to the latter difficulty is to use transplantable tumour lines derived from autochthonous tumours, which typically have a short latency and can be used to rapidly generate preclinical cohorts. Establishment of orthotopic transplantable tumour models with a stable phenotype from GEMM parental tumours would allow therapeutic hypotheses to be tested in a robust, repeatable and timely manner, similar to that of cell line-derived human xenograft models and with the advantage over patient-derived xenografts that the genetic drivers of the model are known in advance. This has been successfully demonstrated in ER negative and BRCA1/2 mouse model systems (Jaspers et al., 2015, Borst et al., 2017, Henneman et al., 2015, Rottenberg et al., 2007) but not yet in a GEM model of ER+ positive breast cancer. Indeed, few such models exist.

Conditional deletion of *Pten* and *p53* in mouse mammary luminal progenitors using Cre recombinase expressed under the control of the *Blg* promoter, gives rise to four main histopathological tumour types. These tumours are representative of at least four different human clinical molecular subtypes of breast cancer, including ER+ luminal tumours (Melchor et al., 2014). However, the dependence of these tumours on ER signalling and their reliance on activation of the PI3K pathway for growth, which would be predicted as a consequence of PTEN loss of function, was never tested. Here, we have taken an orthotopic transplantable tumour approach to establish multiple transplantable lines from two parental *Blg-cre Pten^fl/fl^ p53^fl/fl^* tumours and characterised the impact of tumour passage on the histopathological and genetic profile of these tumours. We found that lines of passaged tumours polarised into mesenchymal-like ER negative tumours and epithelial-like ER positive tumours. The ER positive tumours displayed a range of phenotypes, namely, squamous-, adenomyoepthelial- and adenocarcinoma-type. Signalling pathway dependencies were tested in the epithelial ER positive and mesenchymal-like ER negative tumours. We found that ER positive tumours, but not ER negative tumours, responded to estrogen stimulation; however, the ER positive tumours did not respond to treatment with the specific estrogen receptor downregulator (SERD) fulvestrant. Interestingly, ER positive and ER negative tumours had variable responses to therapy with specific PI3Kα and β inhibitors, in some cases responding to the individual treatments, in others only responding to combination therapy. Our study demonstrates that the transplantable *Blg-cre Pten^fl/fl^ p53^fl/fl^* tumour system is a highly tractable model system for the testing of novel therapeutics and understanding primary resistance.

## Results

### Tumour lines of differing histological subtypes develop from a single adenosquamous carcinoma passaged *in vivo*

We have previously described (Melchor et al., 2014) that in the *Blg-cre Pten^fl/fl^p53^fl/fl^* mouse, mammary tumours of four distinct histotypes develop within the age range of 139-361 days. The four tumour types are classified as adenomyoepithelioma (AME; tumours composed of two distinct neoplastic populations, one being basal-like and one luminal-like), metaplastic adenosquamous carcinomas (ASQC; tumours with similarities to AMEs but with a high percentage of squamous metaplasia and frequent hyperkeratinisation into ‘keratin pearls’), metasplastic spindle cell tumours (MSCT, also known carcinosarcomas; tumours composed predominantly or entirely of fusiform cells which do not express epithelial markers) and adenocarcinoma/invasive ductal carcinomas of no special type (AC/IDC-NST, hereafter simply AC-NST), a histotype with no special features. Notably in this GEMM, the AMEs, ASQCs and AC-NSTs are frequently estrogen receptor positive (Melchor et al., 2014). To examine the plasticity of these phenotypes and their pathway dependencies, we established a series of transplantable lines (‘mouse-derived xenografts’ or MDXs) from frozen fragments of a single parental ASQC tumour which developed in a 147-day old mouse (Figure 1A). This tumour (ID MS1162-1) had a high percentage of cells expressing basal markers (up to 60%) and of squamous metaplasia (approximately 40% of cells having a metaplastic appearance), but also expressed luminal keratins in approximately 10% of cells and ERα in approximately 15% of cells, as we have previously reported (Melchor et al., 2014).

**Figure 1:**
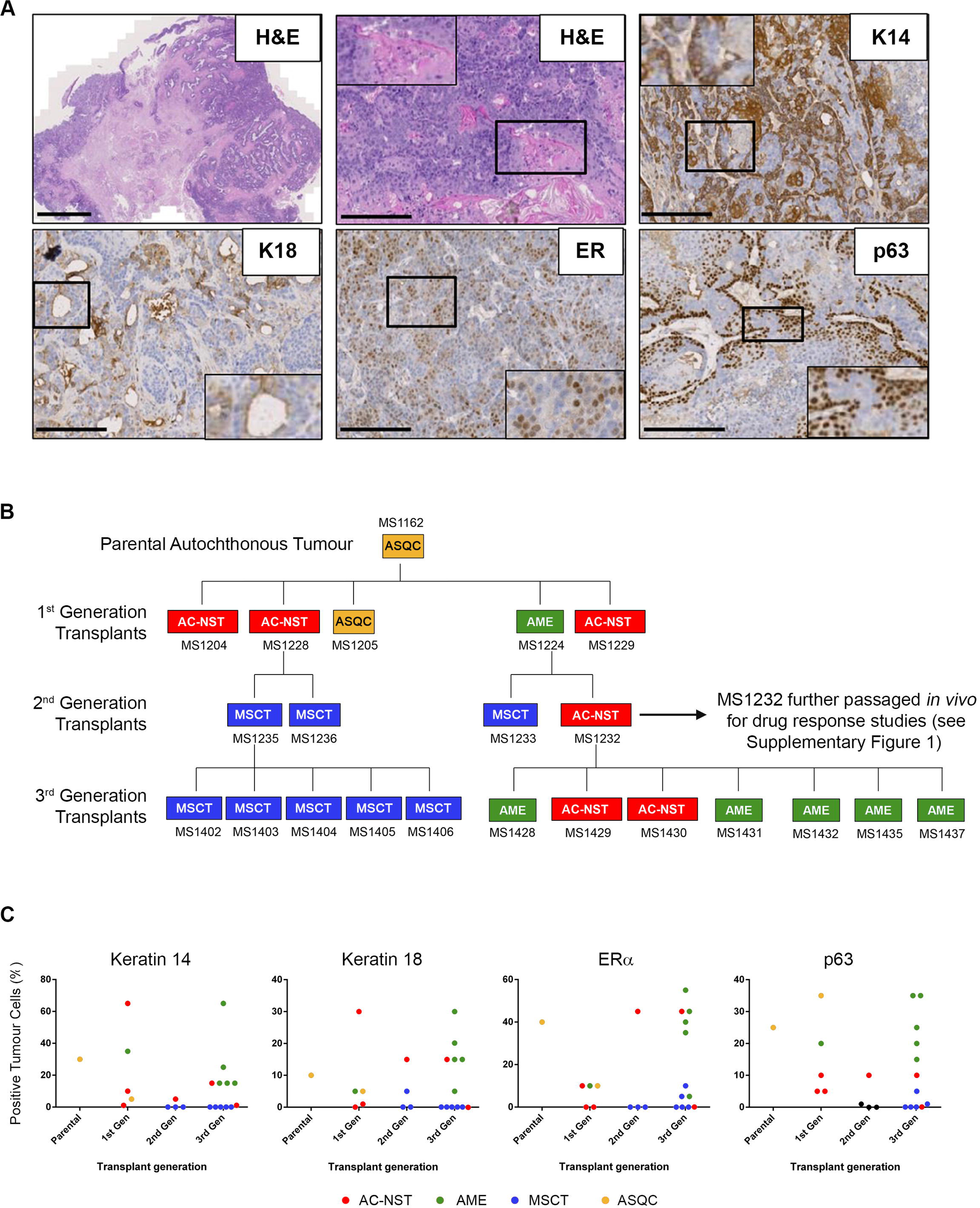
Passage of a single adenosquamous carcinoma gives rise to multiple tumour sub-types. **(A)** Low power H&E section of parental autochthonous metasplastic adenosquamous tumour MS1162; high power H&E showing metaplastic squamous elements (inset) and keratin pearls; Keratin 14 (K14) staining in basal-like cells; Keratin 18 (K18) staining in luminal-like cells; ERα staining in epithelioid cells; p63 staining in epitheloid cells. Scale Bars: (H&E upper left) 2mm; (all other images) 200µm. Insets magnified twice. **(B)** Family tree of parental adenosquamous carcinoma passaged for three generations. Phenotypes of tumours at each generation are indicated and colour-coded (AC-NST, red, adenocarcinoma of no special type; MSCT, blue, metaplastic spindle cell tumour; AME, green, adenomyoepitheliomas; ASQC, yellow metaplastic adenosquamous tumour), together with the reference code for each tumour. Tumour phenotypes were scored as previously described (Melchor et al., 2014, Molyneux et al., 2010). **(C)** Percentages of cells in individual tumours at each generation expressing key histopathological markers (Melchor et al., 2014, Molyneux et al., 2010). Tumour phenotypes are indicated.

Fragments of the parental tumour were transplanted orthotopically into the fourth mammary fat pad of five female nude mice. All five transplants grew palpable tumours. When size limits were reached, mice were humanely killed, tumours were harvested and analysed. Three of these first passage tumours were classed as AC-NSTs with spindle cell elements, one was classed as an AME and one as an ASQC. Full details of the analysis and classification of all tumours within each generation can be found in the supplementary information (Supplementary Table 1). No MSCT tumours developed in the first generation passage but fusiform cells were observed in in MS1228.

One AC-NST (MS1228) and one AME (MS1224) from the first generation were passaged further to give rise to a small cohort of second generation tumours. Interestingly, MS1228 only generated MSCTs, whereas MS1224 gave rise to both an AC-NST and an MSCT. To further assess the phenotypic stability of the transplanted tumours, one second generation MSCT MS1235), and the second generation AC-NST (MS1232) were passaged to establish a third generation cohort (pieces of MS1232 were also frozen and later used to establish cohorts of tumours for testing therapeutic responses; see below). The MSCT gave rise only to daughter MSCTs (5/5), whereas the AC-NST gave rise to both AC-NSTs (2/7) and AMEs (5/7). The full lineage tree is outlined in Figure 1B. All tumours generated in the cohort were high-grade, with high nuclear pleomorphism, little tubule formation and predominantly high mitotic index.

Consistent with our previous work, analysis of staining patterns of our standard panel of immunohistochemical stains (Keratin 14, K14; Keratin 18, K18; p63; ERα) on tumours across the transplant generations showed that MSCTs had low levels of expression of all markers. Expression of K14, K18 and p63 was variable in transplanted AC-NSTs and AMEs, similar to primary tumours (Melchor et al., 2014), with no obvious patterns across the generations. Notably, however, passage resulted in an increase in the proportion of cells in AC-NSTs and AMEs expressing ERα (Figure 1C).

In the three transplant generations, and the primary ASQC, we analysed expression of a panel of genes associated with the three main cell lineages in the normal mouse mammary gland (basal, luminal ER negative and luminal ER+, as well as genes associated with both luminal populations) (Melchor et al., 2014). There were no obvious correlations between gene expression and transplant generation, rather, gene expression generally reflected tumour phenotype irrespective of generation, with MSCTs having low levels of expression of most genes with the exception of *Twist1*, a known regulator of epithelial-mesenchymal transition (Figure 2 and Supplementary Table 2).

**Figure 2:**
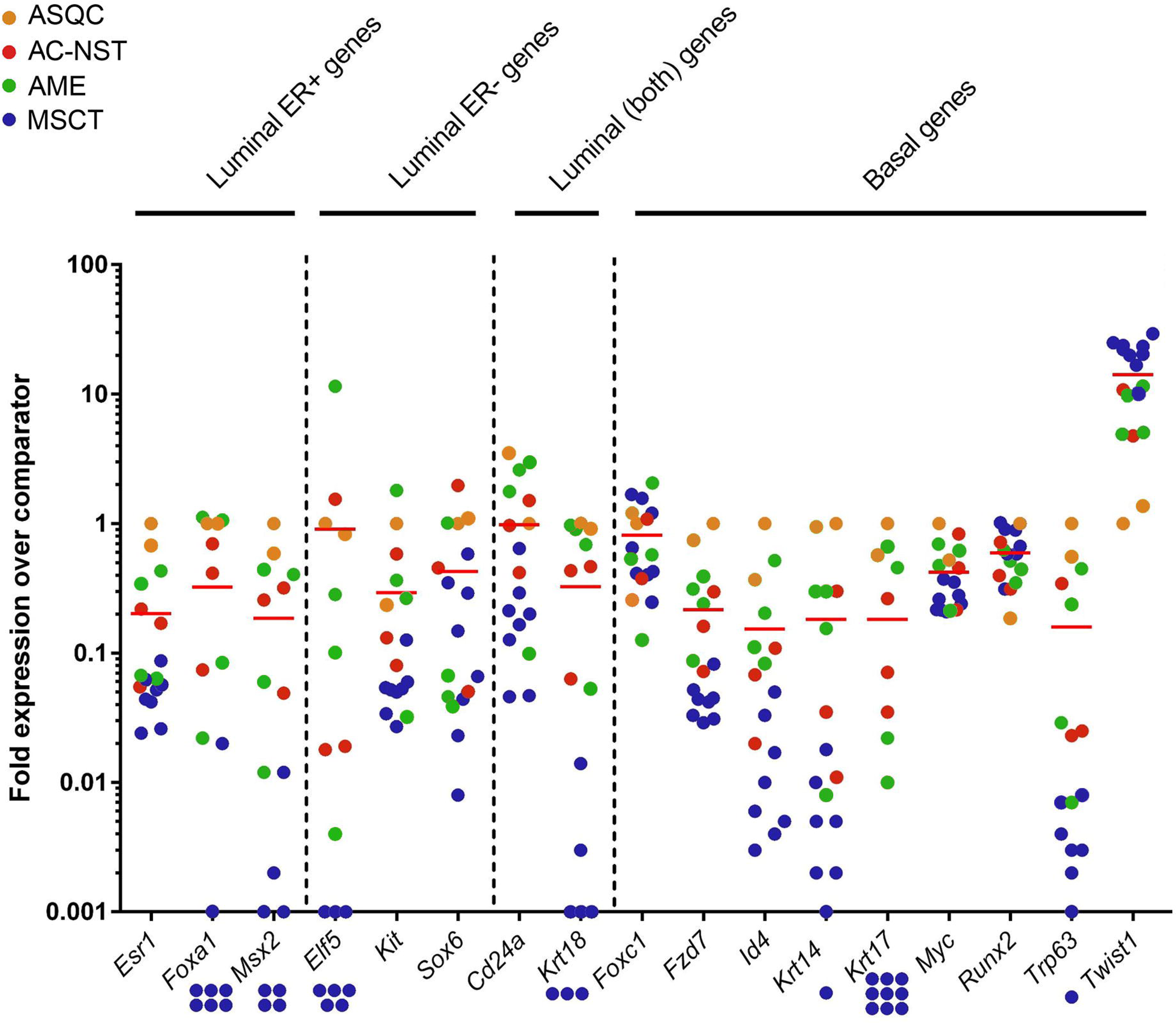
Expression of mammary lineage-associated genes in tumours of different histopathological phenotypes. Genes are divided into those associated with the basal, luminal ER+, luminal ER- or both luminal lineages (Chen et al., 2017). Each point represents expression of a gene in a single tumour, colour coded by phenotype (red, AC-NST; blue, MSCT; green, AME; ASQC, yellow), relative to the expression of that gene in the parental autochthonous ASQC (indicated by the yellow datapoint at ‘1’ for all genes). The mean level of expression within each data set is indicated. Some genes were undetectable in some MSCTs; these are indicated by points below the x-axis.

Therefore, initial heterogeneity of tumours growing from implanted fragments from a single, parental autochthonous tumour developed into tumour lines with either epithelial-like/ERα positive, or mesenchymal-like/ERα negative, features.

### Heterogeneity of response to PI3K pathway inhibition

We next used our transplantable tumour system to examine signalling pathway dependency of the epithelial-like / ERα positive and mesenchymal-like / ERα negative tumour lines. As *Pten* loss is associated with activation of the PI3K pathway and can confer tumour growth dependency on the PI3Kβ isoform in many disease models (Hancox et al., 2015, Schwartz et al., 2015), we focussed specifically on the estrogen receptor (ER) and phosphoinositide 3-kinase (PI3K) pathways, and the effects of their pharmacological blockage on tumour growth.

Frozen fragments of the 2^nd^ generation AC-NST tumour, MS1232, were thawed and used to generate three new generations of tumours. As each generation developed, not only was part of the tumour passaged to start the next generation, but cohorts of mice were implanted with tumour fragments for treatment studies. Furthermore, a new transplantable tumour line was established from a newly developed autochthonous mesenchymal-like tumour, MS1345, which was similarly passaged and used to establish cohorts of mice for treatment (the relationships between these cohorts is shown in Supplementary Figures 1 and 2). Detailed immunohistochemical analysis and phenotyping was carried out on a subset of the tumours derived from these fragments (Supplementary Table 3). Tumours of the MS1345 line retained the MSCT phenotype and were negative for ERα and progesterone receptor A (PRA) (Supplementary Table 3, Supplementary Figure 3A). In contrast to the first set of tumours derived from transplanted pieces of MS1232 (Figure 1), the new set of MS1232-derived tumours were a mix of AC-NST with spindle cell metasplasia and fully metaplastic spindle cell tumours. However, most retained ERα and PRA staining at high levels (indeed, staining was substantially increased compared to MS1232 itself; Supplementary Table 3, Supplementary Figure 3B).

To assess whether passaged tumours derived from the MS1345 and MS1232 lines were estrogen responsive, fragments were implanted into male nude mice with or without supplementation of exogenous estrogen in the form of an estradiol (E2) subcutaneous pellet. The E2 pellet provided a significant growth advantage for tumours derived from the MS1232 ERα/PRA positive line (Figure 3A), whereas E2 supplementation provided no growth advantage to tumours derived from the MSCT MS1345 (Figure 3B).

**Figure 3:**
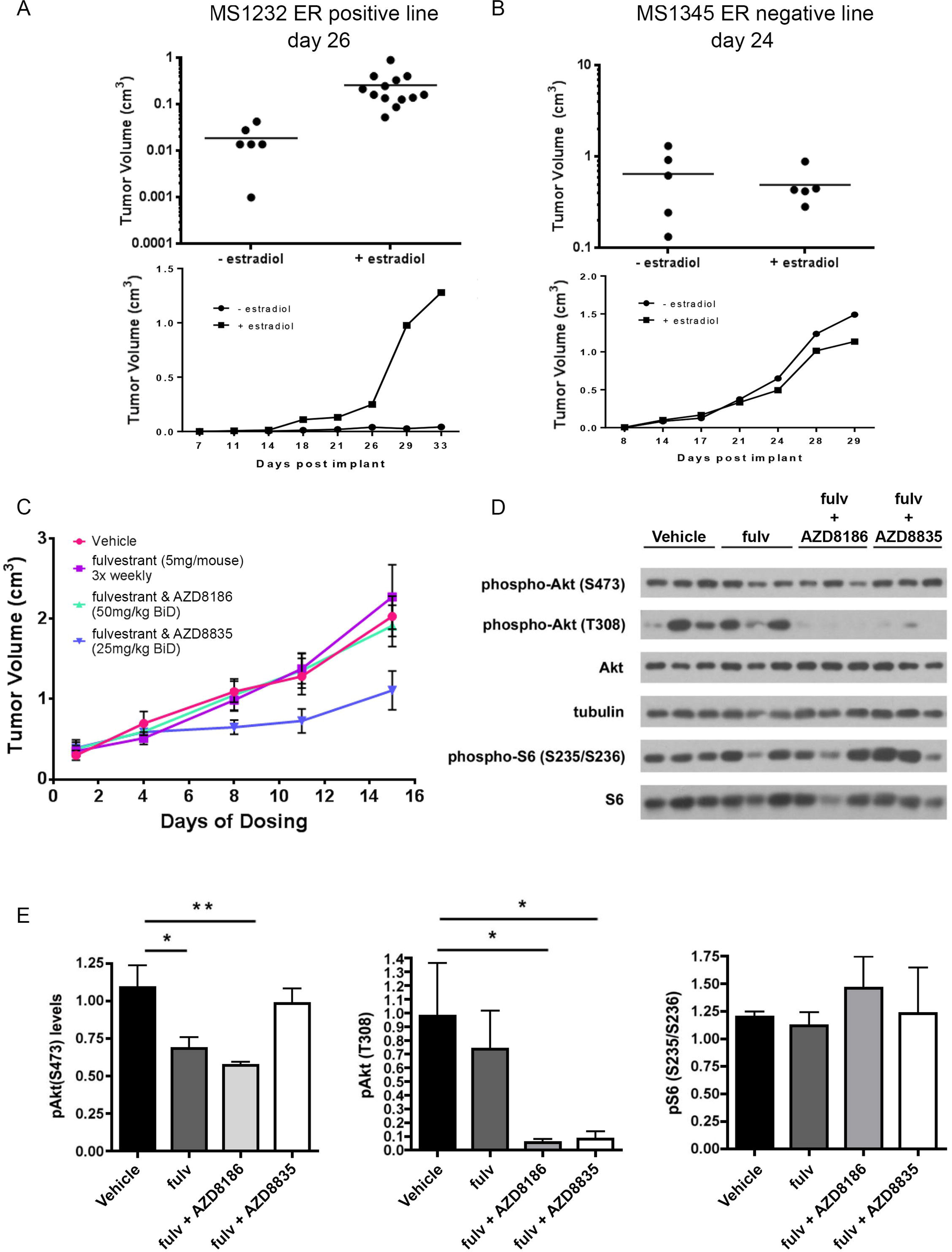
AC-NST tumours require exogenous estradiol for establishment but once established are resistant to selective estrogen degradation therapy. **(A-B)** Growth of tumours from the AC-NST (MS1232) and MSCT (MS1345) lines with and without exogenous estradiol supplementation. **(C)** Treatment of tumours from the AC-NST line with selective estrogen degrader fulvestrant in combination with a PI3K inhibitor (AZD8186) and a PI3Kβ inhibitor (AZD8835). **(D)** Western blot analysis and **(E)** quantitation by densitometry of key downstream markers of PI3K pathway activation in tumours taken at the end of the efficacy study shown in **(C)**. Blots show three representative samples from each group. Densitometry was from a minimum of three, but in some cases four, tumour samples. Unpaired t test. *P<0.05. **P<0.01.

Next, we assessed the response of the estrogen-responsive MS1232 line to a selective estrogen receptor degrader (fulvestrant) with and without inhibition of the PI3K pathway using a PI3Kα specific inhibitor AZD8835 and a PI3Kβ specific inhibitor AZD8186 (Figure 3C). Despite tumours from this line having a significant growth disadvantage when no exogenous estradiol was present, fulvestrant treatment had no impact upon tumour growth in pre-established tumours supplemented with exogenous estradiol (Figure 3C, Supplementary Figure 3C,D). Fulvestrant in combination with either AZD8186 or AZD8835 inhibited phospho-Akt on T308 in tumour samples taken at the end of the efficacy study and a significant reduction in phospho-Akt on S473 was observed in the AZD8186 treatment group (Figure 3D,E). However, only AZD8835 treatment combined with fulvestrant had a significant inhibitory effect on tumour growth (Figure 3C).

We then examined sensitivity of the estrogen insensitive MSCT MS1345 line to PI3K pathway inhibitors using two cohorts expanded from 3^rd^ generation passaged tumours (BLG1402 and BLG1501). When treated with the inhibitors as monotherapies, a modest but significant reduction in tumour size (Figure 4A) was seen in the BLG1402 cohort. Consistent with this, both inhibitors reduced Akt activation in tumours taken at end of study, with AZD8186 appearing to give slightly more suppression of the Akt biomarkers (Figure 4B,C). Tumours treated with AZD8186 also had a reduction in the proximal downstream marker phospho-S6, which was not observed in AZD8835 treated tumours (Figure 4B,C). Unfortunately, insufficient donor tumour material was available to test the combination therapy in this tumour cohort.

**Figure 4:**
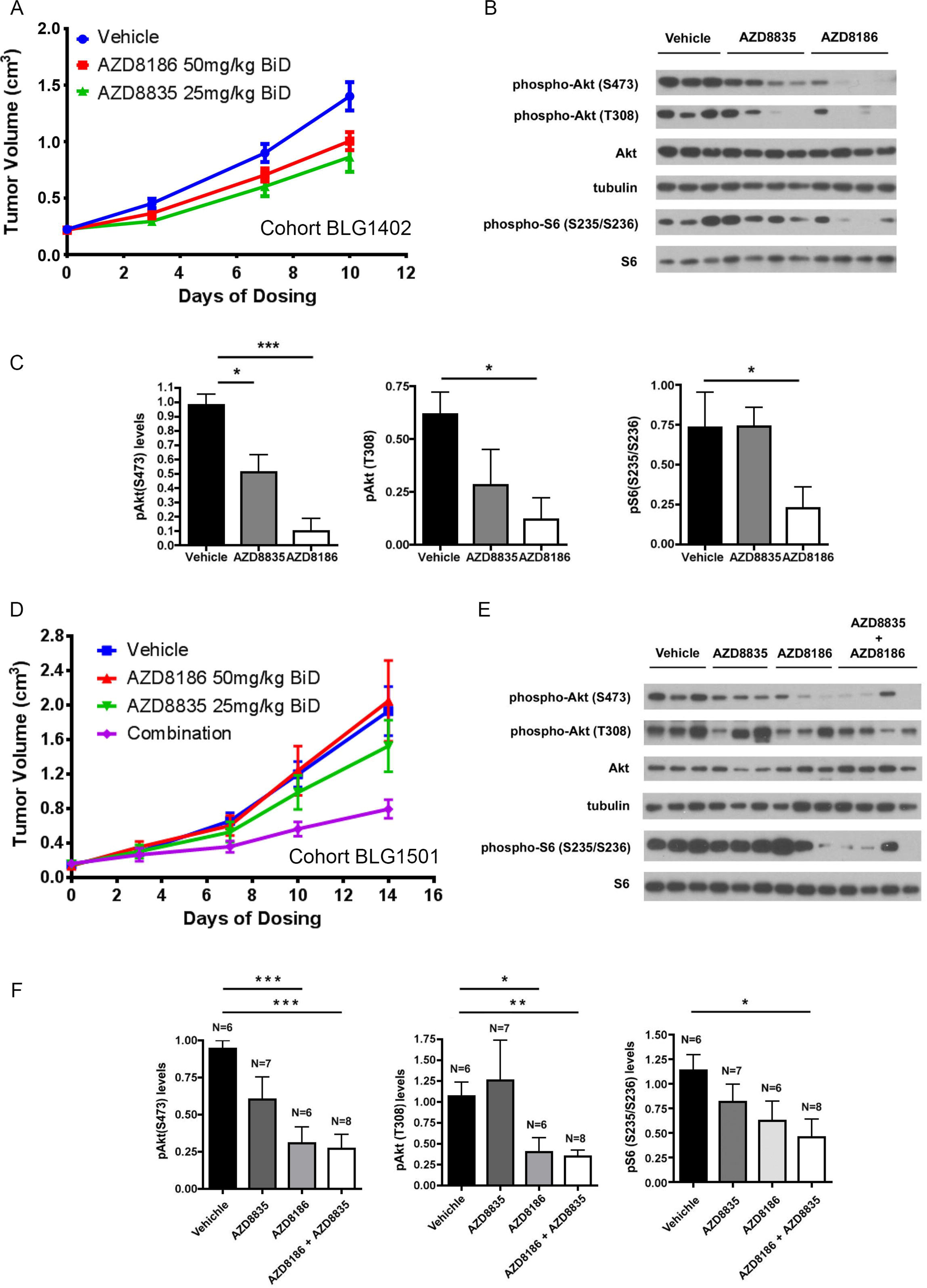
Response to PI3K inhibitors varies in MSCT-derived transplanted tumours established from identical transplant generation. **(A)** Response of MSCT tumours established from BLG1402 (3^rd^ Generation from parental MS1345) fragments to PI3Kα (AZD8835) and PI3Kβ (AZD8186) inhibitors. **(B)** Western blot analysis and **(C)** densitometry of key downstream markers of PI3K pathway activation in tumours taken at the end of study BLG1402 (6h post final dose; n = 4 samples for each group). **(D)** Response of MSCT tumours from BLG1501 (also 3^rd^ Generation from parental MS1345) to PI3Kα (AZD8835)and PI3Kβ (AZD8186) inhibitors. **(E)** Western blot analysis and **(F)** densitometry of key downstream markers of PI3K pathway activation in tumours taken at the end of the study BLG1501 (2h post final dose). Numbers of tumours analysed in each group for densitometry are indicated; blots show at least three representative samples from each group. Unpaired t test. *P<0.05. **P<0.01. ***P<0.001.

In contrast, the BLG1501 cohort was insensitive to single agent treatment with either of the PI3K inhibitors (Figure 4D). However, the combination of both AZD8835 and AZD8186 did reduce tumour growth, suggesting dependency on multiple PI3K isoforms. In end of study samples AZD8835 and AZD8186 both modulated Akt phosphorylation at ser473, though only AZD8186 inhibited phosphorylation at thr308. The combination treatment also inhibited Akt phosphorylation, and delivered the most effective suppression of phospho-S6 (Figure 4E,F). Interestingly, analysis of key biomarkers of PI3K pathway activity in separate cohort of BLG1501-derived tumour samples analysed 2 hours after a single dose of the individual or the combined inhibitors showed that, in contrast to the end-of-study data, there was strong inhibition of pS6 by AZD8835 monotherapy as well as the combined therapies (Figure 5). This suggests that while the PI3K pathway may initially respond to AZD8835 inhibition, it rapidly loses its sensitivity and only the combined inhibitors give a durable response.

**Figure 5:**
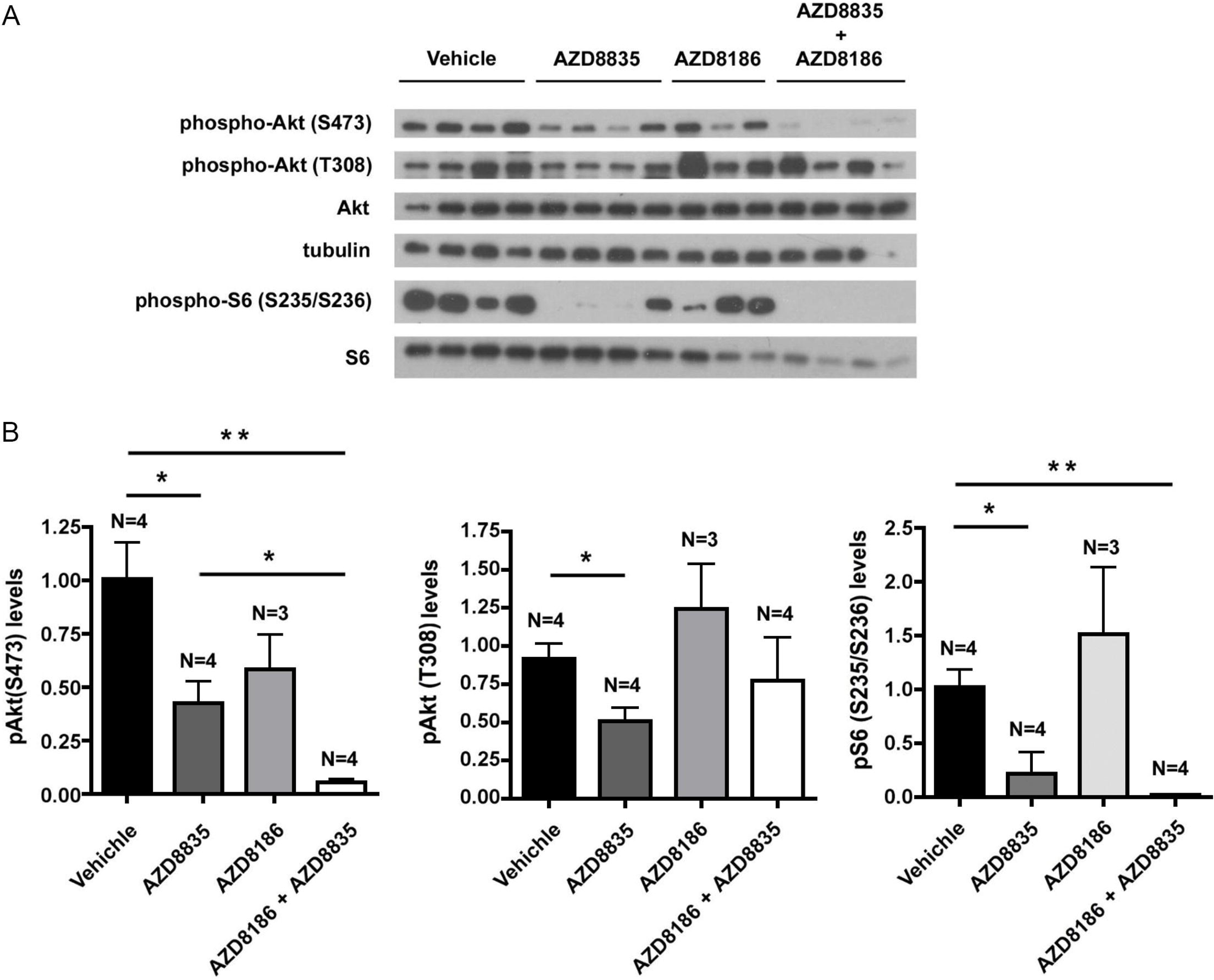
Inhibition of S6 phosphorylation by acute inhibition of PI3Kα but not PI3Kβ. **(A)** Western blot analysis and **(B)** densitometry of key downstream markers of PI3K pathway activation in tumours derived from fragments of BLG1501 treated with only a single dose of each of the inhibitors and samples taken 2h post treatment. Numbers of tumours analysed in each group are indicated. Unpaired t test. *P<0.05. **P<0.01.

Collectively these data suggest that both ERα positive and negative tumours on a *Pten*-null background can require PI3K signalling for growth, and that this can be mediated by both PI3Kα and PI3Kβ isoforms. However, they also show heterogeneity of therapeutic response and differences in pharmacodynamic response in the acute and chronic treatment settings.

### Epithelial-like and mesenchymal-like tumours show distinct mutational profiles

We next asked whether mutational profiles associated with the different lines might explain the response of tumours to therapy. Sixteen whole exomes were sequenced, including the two parental tumours AC-NST MS1232 and MSCT MS1345 and fourteen from tumours representative of the different transplant generations and tumour lines. Of these fourteen, six were from the MS1232 line and eight were from the MS1345 line. One (17_BLG1303) was from an animal which had also been implanted with an estrogen pellet. The remainder were from animals from control, untreated groups from either the estrogen supplementation, fulvestrant or inhibitor treatment groups. The lineage relationships between the tumours used for sequencing is shown in Supplementary Figures 4 and 5. After performing variant calls, we used the COSMIC database list of Cancer Census Genes (Forbes et al., 2016) to focus on genes of potential interest. We looked for mutations in the two parental tumours in genes which might either directly or indirectly affect PI3K signalling, and we looked for new variants in these genes which appeared as the tumours were passaged (Supplementary Table 4). We also looked at mutations in both the parental and passaged tumours in genes not thought to affect PI3K signalling (Supplementary Table 5). We identified between 2900-4400 new variants in the MS1345 line tumours and 1100-2300 in the MS1232 line tumours.

We used two independent methods to assess the relationships between the tumours using the new variants emerging across passages. Hierarchical clustering of these samples based on the common mutations harboured showed a distinct mutational profile of each of the two lines, with two clusters clearly identifying the two lines (Figure 6A). However, two pairs of samples from each of the lines (37_BLG1501 & 45_BLG1501; 55_BLG1409 & 64_BLG1409) did cluster separately from their parental group. This trend was confirmed when running a Principal Component Analysis (PCA) as both pairs clustered separately in the plane defined by the two first principal components, explaining 54% of cumulated variance (Figure 6B). Examination of the mutation patterns in Oncoprint confirmed this observation (Figure 7). These variant samples were both derived from third-generation tumours within their respective transplant lines; however, tumours derived from other third generation tumours did not show such divergence.

**Figure 6:**
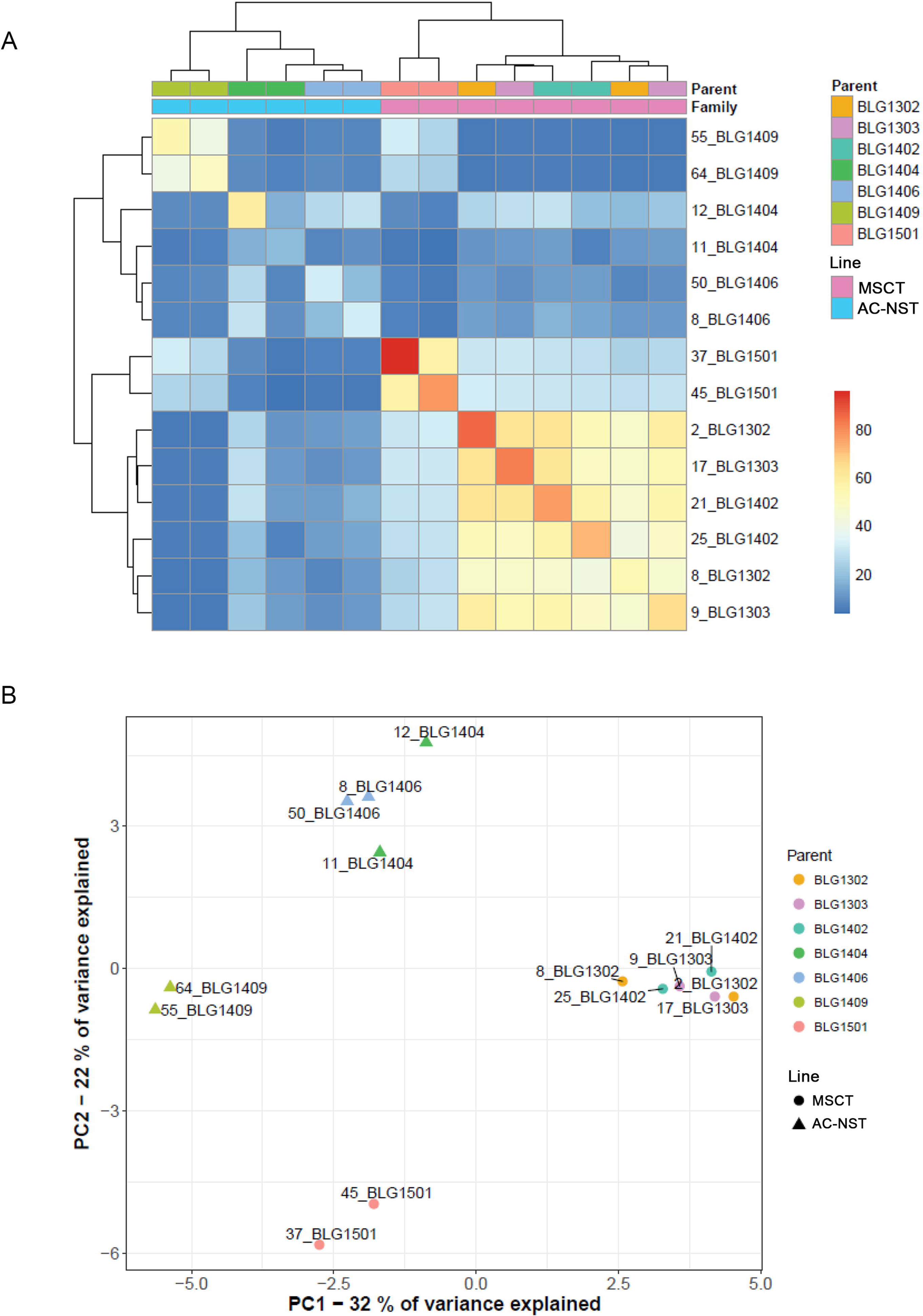
Passaged tumours from distinct lines have distinct mutational profiles. Tumours grown from transplanted fragments of passaged tumours of the AC-NST (MS1232) and MSCT (MS1345) lines underwent exome sequencing and were analysed for new variants occurring in each line during passage. Samples are labelled by cohort e.g. BLG1302 and individual tumour from an animal within that cohort; each cohort was established from a single donor tumour. **(A)** Clustering of exome mutational profiles of tumours derived from transplanted fragments from two independent tumour transplant lines. **(B)** Principal Component Analysis of mutational profiles showing that the profiles of the two tumours from the BLG1409 cohort (numbers 55 and 64; AC-NST line) and the two tumours from the BLG1501 cohort (numbers 37 and 45; MSCT line) were distinct from other tumours from the same transplant lines. These were both third generation tumours within their transplant lines; however, the BLG1406 cohort-derived tumours (AC-NST line) and BLG 1402 cohort-derived tumours (MSCT line) were also third generation, therefore, passage number alone cannot explain this divergence.

**Figure 7:**
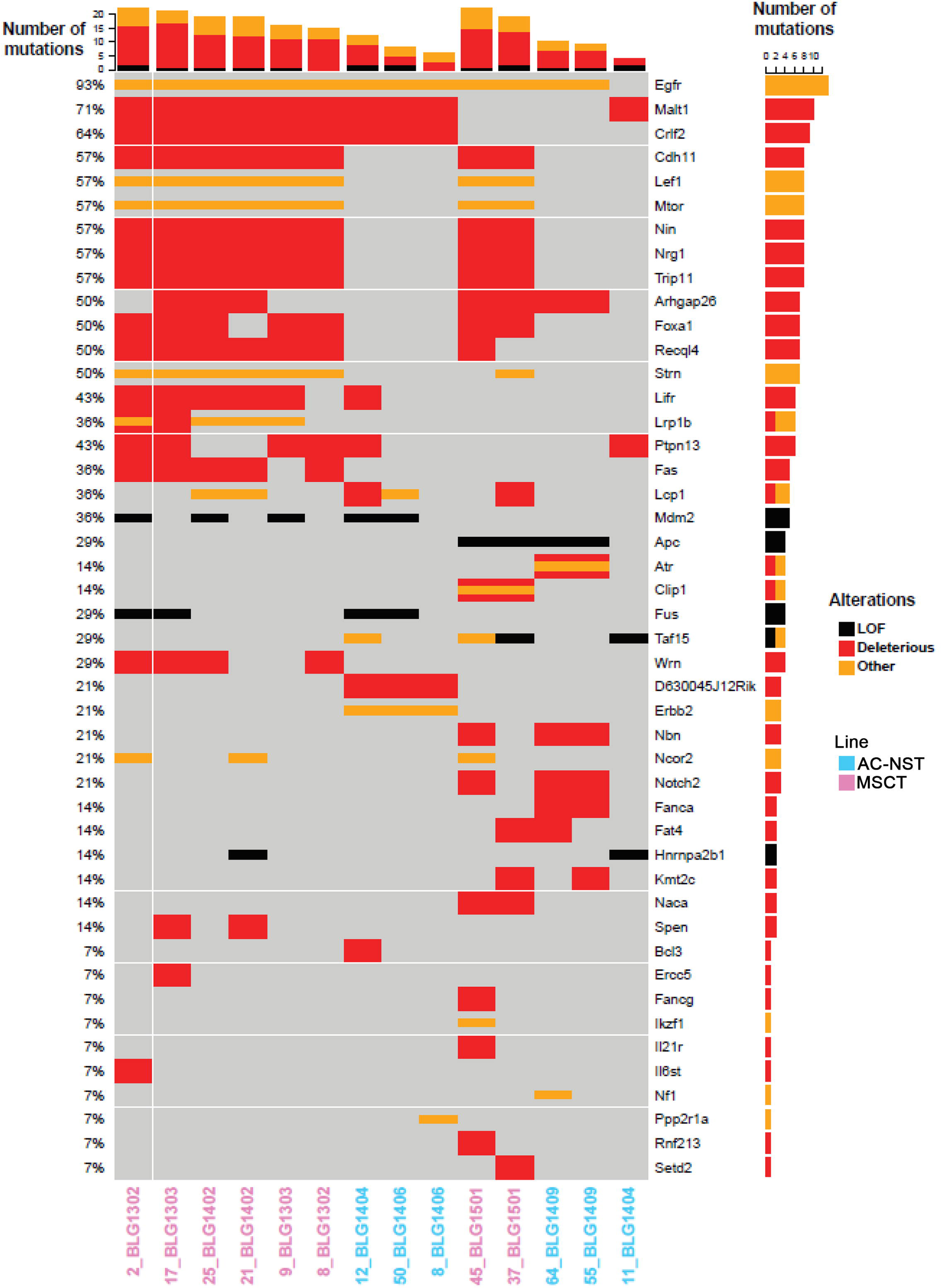
Passaged tumours from distinct lines have both distinct and common variants. Oncoprint analysis of new variants arising in the AC-NST (MS1232) and MSCT (MS1345) lines during passage. Functional consequences of the variants are indicated. Sample numbers are colour-coded to indicate origin from the epithelial (blue) or mesenchymal (pink) origin line.

Mutations were identified which were unique to individual tumours (e.g. R359Q BCL3 in 12_BLG1404), which were present in multiple tumours from the same line across generations (e.g. H389R FOXA1 in the MSCT MS1345 line and P126L MALT1 in AC-NST MS1232) and, notably, some which were identified in tumours from the two different lines (e.g. N1980del APC; L22F ARHGAP26; G575G EGFR). Given that the two lines of passaged tumours used for the exome analysis were not related, this may suggest selection for mutational events which confer a growth advantage during passage. Other remarkable mutation patterns suggest the selective pressures of transplantation drive convergent evolution and the emergence of clones in which particular mutations have been strongly selected for. For example, in the AC-NST MS1232 line, two different mutations in LCP1, a G99E mutation and a C>T mutation at S418, which does not alter the coding sequence but may affect a splice site, appear in tumours 12_BLG1404 and 50_BLG1406 respectively. In the independently derived MSCT MS1345 line, LCP1 mutations also appear in three tumours, 21_BLG1402 and 25_BLG1402, both of which also carry the C>T splice site mutation, and a T527M mutation in 37_BLG1501. LCP1, also known as L-Plastin, is an actin-binding protein which is reported to be regulated by a PI3K – AP4 signalling axis (Chen et al., 2017).

No mutations were identified in either *Esr1* or *Pik3ca* which might have affected response to fulvestrant or PI3K inhibitors. Mutations in *Pik3cb* were detected in one tumour from the MSCT BLG1501 cohort (#37; highlighted in Supplementary Table 4), although it is not obvious how this would explain the lack of sensitivity of tumours in this cohort to AZD8835 and AZD8186 monotherapies but not to the combination. A mutation in the ERα pioneer factor (Ross-Innes et al., 2012) *Foxa1* was detected in the AC-NST ER positive founder tumour, MS1232, but this was apparently lost (presumably due to clonal selection) as the tumours were passaged, while an identical mutation was selected for in the unrelated MSCT ER negative line as the tumours were passaged. Therefore, changes in mutational profile in the passaged tumour lines likely reflect selective pressures associated with transplantation but do not obviously explain differences in response to treatment.

## Discussion

Use of transplantable tumours derived from more indolent GEMMs is an attractive strategy to increase the size and number of tumour studies that can be run over time. Increasingly there is a desire to use these models to support testing of novel therapeutics, targeting both the tumour and immune cells. Therefore, being able to establish transplantable tumours from these models is essential given the time limitations and cost of generating autochthonous tumour bearing animals. The autochthonous *Blg-cre Pten^fl/fl^p53^fl/fl^* mouse model is known to develop a diverse range of histologically defined tumour types (Melchor et al., 2014). In our current study, a single tumour of adenosquamous phenotype from one of these animals is able to give rise to four distinct tumour phenotypes. The phenotype of the tumours assessed was more diverse in the initial generations, but this variability decreased with passage/generation. By the second generation, the tumour phenotypes that developed could be generally classified into two distinct groups, one being a more homogenous set of ER negative spindle cell tumours, the other being more heterogeneous but broadly classifiable as epithelial and ER positive.

Tracking the generational evolution of these tumours has given an interesting insight into the heterogeneity of the tumour subtypes present in this model. The epithelial phenotypes varied between ASQC, AME and AC-NST. It is important to note that ASQC and AME tumours are very similar in appearance and gene expression (Melchor et al., 2014) and in this study we could observe small numbers of metaplastic squamous cells in tumours we classed as AME and AC-NST (20% of cells showing metaplastic features was considered as the cut-off point at which an AME with squamous elements would be classified as an ASQC). It is likely that a combination of potential selective pressures of growth and transplantation could alter the proportion of luminal-like, basal-like and squamous cells that form the basis of tumour phenotyping. In addition, the likely sampling bias associated with assessment of a single section from one piece of a tumour should be taken into account. Therefore, it reasonable to consider that the epithelial, ER positive, AME/ASQC/AC-NST tumours should be considered as a single group of epithelial phenotype as opposed to the spindle cell tumours of mesenchymal phenotype. Importantly, however, the epithelial tumours retained ER expression whereas the mesenchymal tumours lost ER. One exception to this was the set of tumours derived from frozen pieces of the epithelial ER+ tumour MS1232, used for drug response studies, which acquired spindle cell metaplastic features, or underwent full metaplasia (or, indeed, the epithelial cells were overgrown by small numbers of spindle cells which had not been detected in the piece of MS1232 which was analysed), yet retained ER expression.

To explore whether diversity of tumour phenotype was also reflected in pathway dependency, we assessed how modulation of the ER and PI3K pathways impacted on tumour growth in two distinct tumour phenotypes. The ER+ epithelial tumours were dependent on exogenous estradiol for tumour growth. However, they did not respond to single agent treatment with the selective estrogen degrader fulvestrant, but were responsive to combination therapy with the PI3Kα inhibitor. This is consistent with reports that activation of PI3K signalling confers resistance to endocrine therapy in ER+ preclinical breast cancer models (Toska et al., 2017). We cannot exclude the possibility that high levels of estradiol present in these animals may contribute to a lack of response of fulvestrant. However, response to fulvestrant is observed in other estradiol supplemented ER+ preclinical tumour models (Weir et al., 2016).

In contrast the ER-model was not dependent on estradiol for growth, consistent with its lack of PR expression. Consistent with loss of PTEN, this model responded to PI3K pathway inhibition. However, the dependency on an individual PI3K isoform varied between experiments performed on different cohorts established from the same tumour line. Loss of PTEN is known to confer dependency on the PI3K pathway and it is thought that PTEN loss is largely dependent on PI3Kβ (Wee et al., 2008, Ni et al., 2012). Cohort BLG1402 tumours (derived from BLG1303 tumour #19), were sensitive to single agent treatment of either inhibition of PI3Kα or PI3Kβ, but cohort BLG1501 tumours (derived from BLG1303 tumour #18), were only responsive to dual inhibition. It is known that reciprocal feedback reactivation of PI3Kα or β can occur in PTEN null tumours requiring inhibition of both isoforms to deliver maximum efficacy (Schwartz et al., 2015). This suggests that Pten null tumours derived from the *Blg-cre Pten^fl/fl^p53^fl/fl^* tumour fragments may be driven by both PI3K isoforms, and the dominant isoform may vary between donor tumour fragments. Evidence to support this is seen in the end of study biomarkers analysis, where although PI3K biomarkers are modulated upon treatment, the pattern of biomarker modulation to compound treatment is different between cohorts.

Our findings underline the potential for the confounding effects of tumour phenotype instability on results based on serially passaged mouse tumour xenografts. However, while variability of response to treatment was seen between transplanted tumour cohorts in different experiments, within any one study the response of tumours in any study arm was remarkably consistent. This reinforces the importance of establishing a sufficiently powered cohort for a treatment study based on transplant of a single donor tumour and the requirement to look at multiple cohorts established from multiple donors. Variability with passage is more profound in mouse xenografts than human patient xenografts. In a recent study, the histological and molecular phenotypes, including the intra-tumour heterogeneity, of 23 patient-derived breast cancer xenografts were found to be remarkably stable (Bruna et al., 2016). While the original drivers of such tumours cannot be determined, around 20% of the ER-tumour xenografts and 30% of ER+ xenografts in this study were found to carry *PIK3CA* mutations. Interestingly, use of cells derived from these xenografts as models for high throughput drug screening demonstrated the *PI3KCA* mutant tumours responded to PI3Kα and AKT inhibitors, but not mTOR and PI3Kβ inhibitors. PTEN mutations were not reported in this PDX dataset (Bruna et al., 2016).

Exome sequencing did not reveal a common pathway or node that would confer resistance to fulvestrant or PI3K isoform dependencies, although mutations in FOXA1 were suggestive. FOXA1-mediated reprogramming of ERα binding to DNA has been linked to therapy resistance in ER+ breast cancer (Ross-Innes et al., 2012) and whether FOXA1 has a role in the mechanism of fulvestrant resistance in our model system warrants further investigation. However, a general accumulation of new mutations and recurrent mutations in genes from transplanted tumour lines with independent origins suggests such mutations may be important for surviving transplantation and arguably, by extension, metastasis. The actin-binding protein LCP1, or L-plastin, is regulated by PI3K signalling, and is linked to metastasis in both prostate and breast cancer (Chen et al., 2017, Inaguma et al., 2015). Furthermore, plastin polymorphisms predict colon cancer recurrence after therapy (Ning et al., 2014) and plastin3 (or T-plastin) identifies circulating tumour cells undergoing EMT and predicts survival in colorectal and breast cancer (Ueo et al., 2015, Yokobori et al., 2013). The role of LCP1 in tumour invasion and migration remains to be fully understood, as does its regulation by PI3K signalling, but the presence of identical mutations in independent tumour lines following transplantation suggest that it – or the functional pathway in which it operates – is a key player in surviving the selective pressures associated with movement to, and growth in, a new tissue site.

Therapeutic testing has been carried out in autochthonous breast models similar to the *Blg-cre Pten^fl/fl^p53^fl/fl^* model (Karim et al., 2013, Hay et al., 2009). These experiments are time-consuming and costly, and due to the heterogeneous nature of these models, animals will be recruited onto treatment without prior knowledge of its tumour phenotype and mutational landscape (aside from its driver mutations). By using a transplantable version of these models, each experimental cohort represents a unique variant of the autochthonous model. Therefore, therapeutic testing of multiple cohorts using this approach builds understanding of the diversity of response to therapeutics in each tumour subtype. Clearly, careful characterisation of the features of each cohort is required to interpret data appropriately. Despite this, the transplantable model we present here provides an opportunity to examine response to therapies in a tumour of defined pathology and genetics, in a timescale conducive to high-throughput testing similar to human xenograft models.

## Materials and Methods

### *In vivo* experiments

Animal experiments were carried out at both Cardiff University and AstraZeneca under the authority of Home Office project and personal licences. Experiments were compliant with the UK Animals (Scientific Procedures) Act, which is consistent with EU Directive 2010/63/EU, conformed to ARRIVE guidelines and had undergone internal ethical review. Tumour fragments were implanted subcutaneously via trochar in the 4^th^ inguinal mammary fat pad of male and female athymic nude mice (Athymic Nude-*Foxn1^nu^*, Envigo, Huntingdon, UK). Some male mice were implanted with a subcutaneous 0.5 mg/21 day 17β estradiol pellet (Innovative Research of America, Sarasota, Florida, USA) one day prior tumour fragment implant. Tumour growth was monitored by calculating tumour volumes via calliper measure using the equation: 3.14 x length x width2/6000.

### Tumour Phenotyping

For tumour phenotyping, animals were euthanised when tumour sizes reached specified endpoints. Tumours were excised under aseptic conditions. One piece was minced to 1mm^3^ fragments which were frozen in FBS supplemented with 10% DMSO, one piece was snap-frozen for DNA, RNA and protein extraction, and one piece was fixed in either 4% paraformaldehyde or 10% neutral buffered formalin, processed by standard methods and embedded in paraffin wax for histopathological characterisation. Tumours were phenotyped from H&E sections and immunohistochemical staining according to our previous criteria (Melchor et al., 2014).

### Drug Treatment Studies

To test the dependency of the tumours on the ER and PI3K pathways animals were recruited onto study once tumours had reached a volume of 0.2-0.35cm^3^. Animals were treated with the following agents: selective estrogen receptor degrader (SERD) fulvestrant: 1mg/mouse, subcutaneous three times weekly, formulated in peanut oil; AZD8835: oral twice daily, formulated in HPMC/Tween; AZD8186: oral twice daily, formulated in HPMC/Tween. Animals were treated for up to three weeks and tumour volume monitored over this period. Statistical analysis of tumour volumes was carried out as previously described (Shaw et al., 2017).

### Immunohistochemistry

Tissues were processed and immunohistochemistry carried out as previously described either manually (Molyneux et al., 2010, Robertson et al., 2008) or using autostainers (Davies et al., 2015). Primary antibodies used were as follows: cytokeratin 14 (clone LL002, 1:500, #ab7800, Abcam, Cambridge, UK), cytokeratin 18 (clone Ks18.04, 1:5, #65028, Progen), Estrogen Receptor (clone 6F11, 1:500, #VP-E613, Vector Labs, Peterborough, UK), Progesterone Receptor (clone hPRa7, #MA5-12658, Thermo Scientific, Loughborough, UK), p63 (clone BC4A4, 1:100, #ab735, Abcam), Ki67 antigen (clone MM1, 1:100, VP-K452, Vector Labs,).

### RNA extraction and Gene Expression analysis

RNA was extracted from 10-15mg of frozen tumour tissue using the RNeasy Plus Kit according to the manufacturer’s protocol (Qiagen). cDNA synthesis was carried out using QuantiTect Reverse Transcription Kit (Qiagen) according to the manufacturer’s protocol. Taqman Assays-on-Demand probes were used to perform gene expression analysis on the QuantStudio Flex Software detection system. Probes used were can be found in Supplementary Table 6. Actb (β-actin) and B2m (β2-microglobulin) and Ubc (Polyubiquitin-C) were used as endogenous controls and results calculated using the ΔΔC_t_ method.

### Protein extraction and Western blot analysis

Tumours were homogenized in Cell Lysis buffer (#9803, Cell Signalling Technology, NEB, Hertfordshire, UK) with 1mM PMSF in Lysing Matrix D tubes (MP Biomedicals, Fisher Scientific, Loughborough, UK) on a Precellys 24 machine (Bertin Instruments, Labtech International, Heathfield, UK) and passed through a 23G needle prior to incubation on ice for 30 minutes. Total cell extracts were then centrifuged at 10000g at 4°C and supernatants were assessed for protein concentration using the BCA assay kit (Pierce, Thermo Scientific) according to the manufacturer instructions. Protein samples (20 micrograms/lane) were separated by SDS-PAGE, transferred to PVDF membranes (IPVH00010, Merck Millipore, Hertfordshire, UK) and immunoblotted with antibodies to phospho-S473 Akt (rabbit polyclonal, #9271, Cell Signalling Technology), phospho-Thr308 Akt (rabbit monoclonal, clone 244F9, #4056, Cell Signalling Technology), phospho-Ser235/236 S6 Ribosomal Protein (rabbit monoclonal, clone D57.2.2E, #4858, Cell Signalling Technology), Akt (rabbit monoclonal, clone 11E7, #4685, Cell Signalling Technology), S6 Ribosomal Protein (rabbit monoclonal, clone 5G10, #2217, Cell Signalling Technologies). Alpha-tubulin (mouse monoclonal, clone DM1A, #T9026, Sigma, Poole, Dorset, UK) was used as loading control. Resulting immunocomplexes were detected by HRP-conjugated anti-mouse IgG (A4416, Sigma) or anti-rabbit IgG (A6154, Sigma) secondary antibodies and enhanced chemiluminescent (ECL) reagents (WBLUF0100, Merck Millipore).

### DNA extraction and Exome sequencing

Genomic DNA was extracted from either frozen tumour tissue using the QIAamp DNA Mini Kit (Qiagen) or from FFPE tissues (tumour MS1345) using the QIAamp DNA FFPE Tissue Kit (Qiagen) according to the manufacturer’s protocol and DNA quality tested by running on an agarose gel. Exome sequencing was carried out externally at Perkin Elmer according to their in-house protocol, 100bp reads using Illumina HiSeq instruments.

### Sequence alignment and variant calls

The raw fastq files were processed using bcbio (https://github.com/chapmanb/bcbio-nextgen). Within bcbio, bwa 0.7.12 (Li and Durbin, 2010) was used for sequence alignment, samblaster 0.1.22 (Faust GG, Hall IM. 2014. SAMBLASTER: fast duplicate marking and structural variant read extraction. Bioinformatics 30:2503–2505) to mark duplicates, VarDict Java 1.4.8 (Lai et al., 2016) to call variants in paired variant calling mode and SnpEff 4.3g (Cingolani et al., 2012) to annotate effects.

### Variant filtering and annotation

In all subsequent analysis, variant calls impacting canonical coding transcripts were focused on, where canonical transcripts were defined as the transcript with the longest CDS for each annotated gene. Any synonymous variant was excluded (functional class “synonymous_variant”) and germline variants were filtered out using data from the Mouse Genome project (strain C57BL/6). Finally, in order to increase confidence in our calls, variants with less than 10 supporting reads (estimated by multiplication of Allele Frequency by depth) were not considered. Variants with functional classes (SNPEff annotation) “frameshift_variant”, “stop_gained”, “disruptive_in_frame_deletion” and “disruptive_inframe_insertion” were predicted to result in loss of function. We then used SIFT (Ng and Henikoff, 2003) to classify missense variants and distinguish between tolerated and deleterious ones. The list of cancer census genes and their annotation was obtained from the COSMIC database (Forbes et al., 2016).

### Sample clustering and results representation

New variants identified in the passaged tumours were referenced using a unique key composed of chromosomal coordinates and nucleic acid changes. Each sample was then represented by a vector describing the presence of these variants in the samples, the value 1 indicating successful detection, 0 absence of the variant. Hierarchical clustering based on Euclidean distances was then performed on these vectors. Clustering results and heatmap were displayed using the pheatmap package (https://cran.r-project.org/package=pheatmap). Principal components analysis was also run using these numerical values. The “Oncoprint” visualisation depicting sample and gene clustering was generated using the ComplexHeatmap (Gu et al., 2016) Bioconductor package.

## Acknowledgements

The authors would like to acknowledge all members of the EU IMI PREDECT consortium, Amar Rahi and AstraZeneca Laboratory Animal Sciences group for assisting with *in vivo* experiments and Emma Nuttall for tissue processing support.

## Data Availability

NGS data has been deposited in The European Nucleotide Archive (www.ebi.ac.uk/ena): submission reference number PRJEB31181

## Author Contributions

HJD and HM jointly carried out experimental work, collected, analysed and interpreted data and participated in the writing of the manuscript. GT, HK, CC, MJA, SL carried out experimental work, collected and analysed data and contributed to the manuscript. MJS and STB planned and supervised the work, obtained funding, analysed and interpreted data and wrote the manuscript. All authors reviewed and approved the final manuscript.

## Supplementary Information

### Supplementary Figure Legends

**Supplementary Figure 1:** Generational relationships of AC-NST ER+ transplanted tumour cohorts used for estrogen, fulvestrant and PI3K inhibitor response experiments.

**Supplementary Figure 2:** Generational relationships of MSCT ER-transplanted tumour cohorts used for PI3K inhibitor response experiments.

**Supplementary Figure 3: (A)** Representative images of H&E and PR staining of three tumours from the autochthonous MSCT MS1345 line, including the parental tumour and representatives of the 1^st^ and 2^nd^ transplant generations. Scale bars = 100µm. **(B)** Representative images of H&E and PR staining of three tumours from the AC-NST MS1232 line, including the parental, 1^st^ and 2^nd^ generations. Scale bars represent 100µm. **(C)** Response of AC-NST MS1232 transplant line-derived tumours to fulvestrant grown in female nude mice and **(D)** representative PR staining of two control and two fulvestrant-treated tumours from these groups. Scale bars = 250 µm.

**Supplementary Figure 4:** Lineage relationships of AC-NST ER+ transplanted tumours used for exome sequencing. Solid ovals indicate tumours which were sequenced; the lightly shaded oval (1_BLG1401) indicates a tumour from which material was not available for sequencing but which was used to propagate subsequent generations.

**Supplementary Figure 5:** Lineage relationships of MSCT ER-transplanted tumours used for exome sequencing. Solid ovals indicate tumours which were sequenced; lightly shaded ovals indicate tumours from which material was not available for sequencing but which was used to propagate a subsequent generation.

### Supplementary Tables

**Supplementary Table 1: Features of parental autochthonous and passaged tumours.** Tumours were scored as described (Melchor et al., 2014, Molyneux et al., 2010). The presence of cells with metaplastic (spindle or squamous) features was considered first (with a cut-off of 20% considered to be sufficient to diagnose a tumour as MSCT or ASQC respectively). For tumours which did not fall into either of these categories, the presence of distinct basal-like and luminal-like neoplastic cell populations (defined by proportion and pattern of p63 staining) was used to determine whether the tumour was an AME or AC-NST.

**Supplementary Table 2: Gene expression levels relative to comparator parental tumour for tumours depicted in** **Figure 2**.

**Supplementary Table 3: Characterisation of tumours derived from implanted fragments used for PI3K pathway inhibition studies.**

**Supplementary Table 4: Variants detected in parental and passaged tumours reported to impact on PI3K signalling.**

**Supplementary Table 5: Variants detected in parental and passaged tumours not related to PI3K signalling.**

**Supplementary Table 6: TAQman probes used for qrtPCR studies.**

