## Supplementary figures and images for "Generation and characterisation of an estrogen receptor-positive GEMM-derived *Pten p53* null transplantable breast tumour model for therapeutic testing"

### Supplemental Figure S1

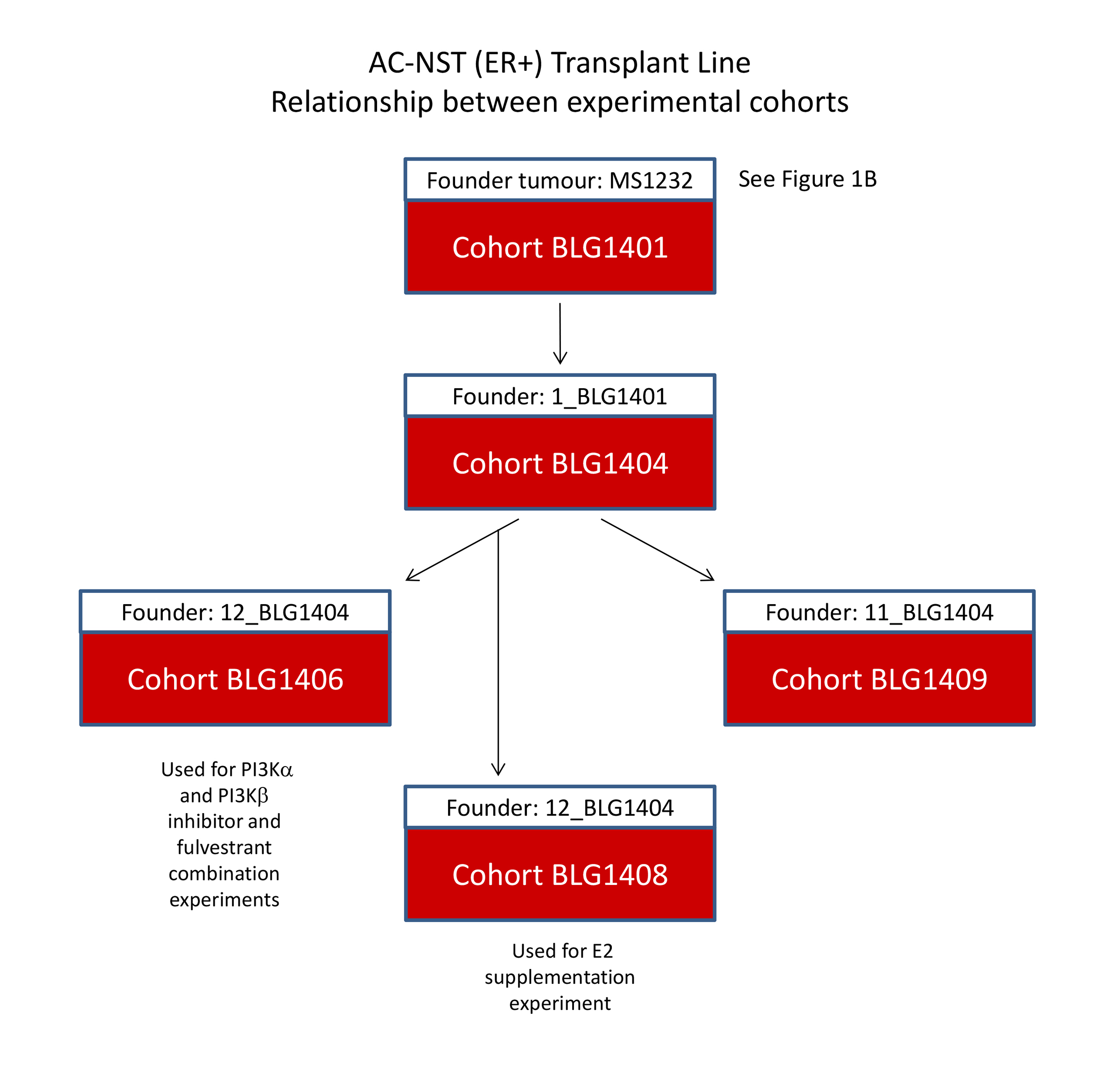

### Supplemental Figure S2

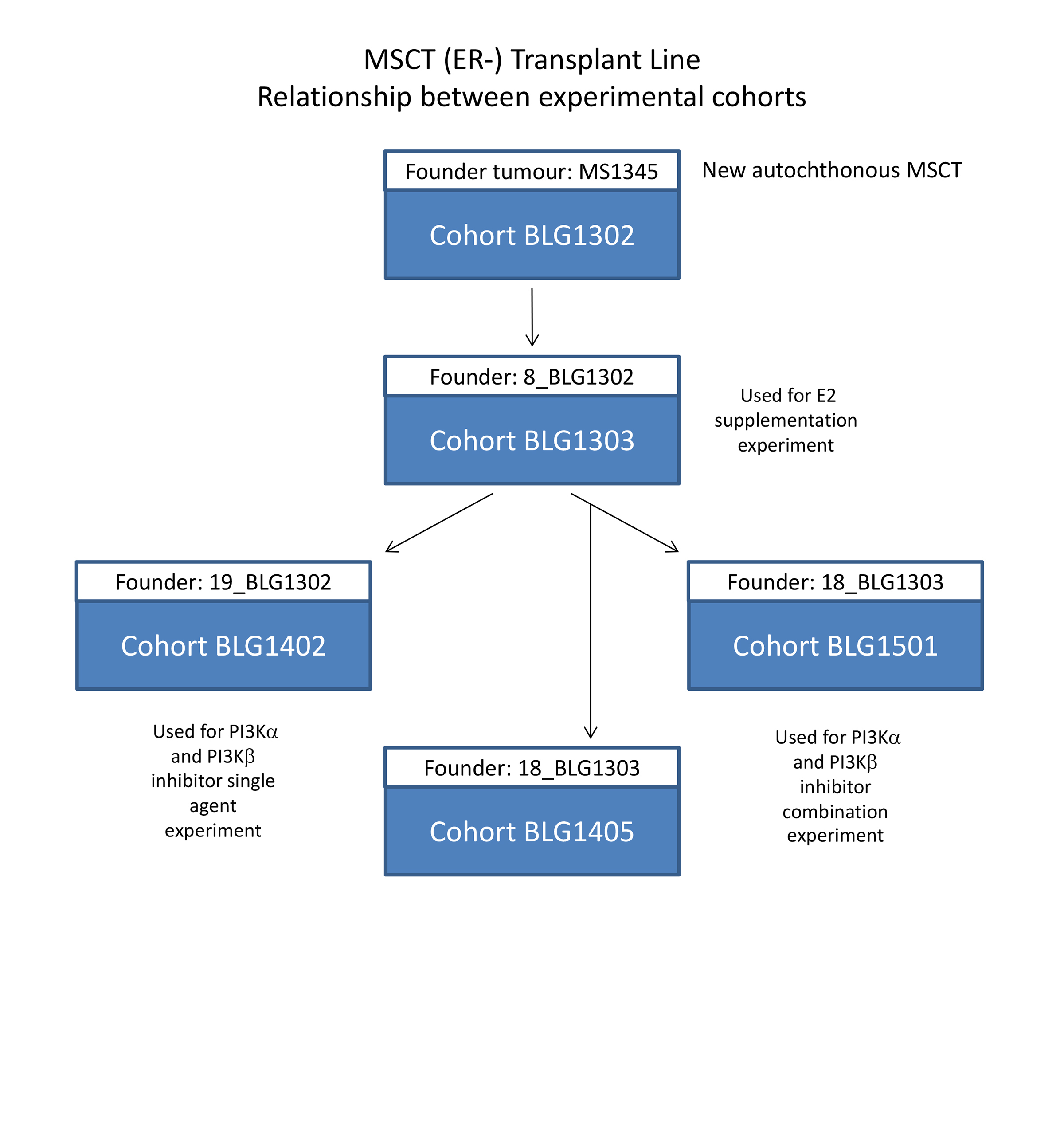

### Supplemental Figure S3

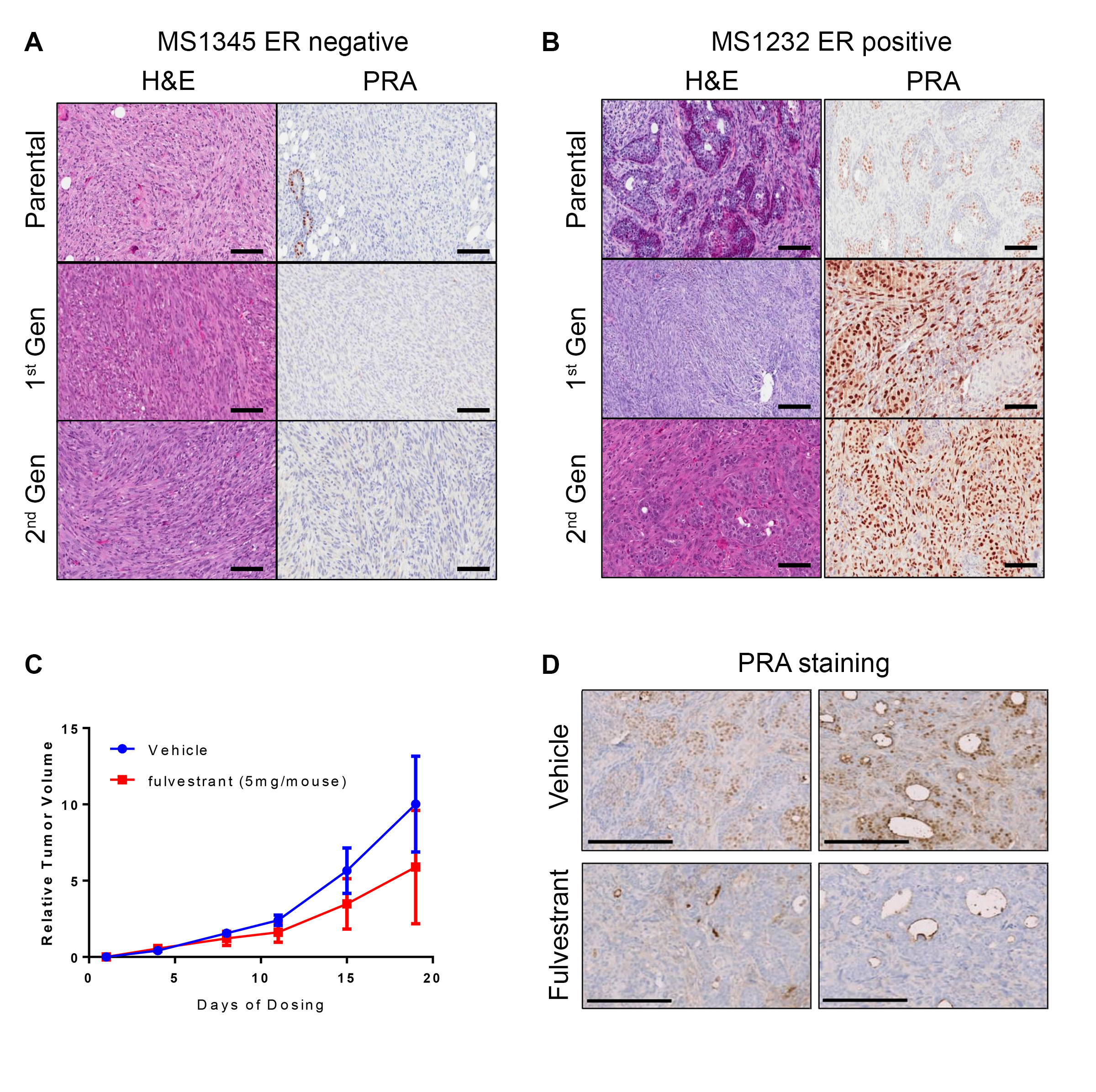

### Supplemental Figure S4

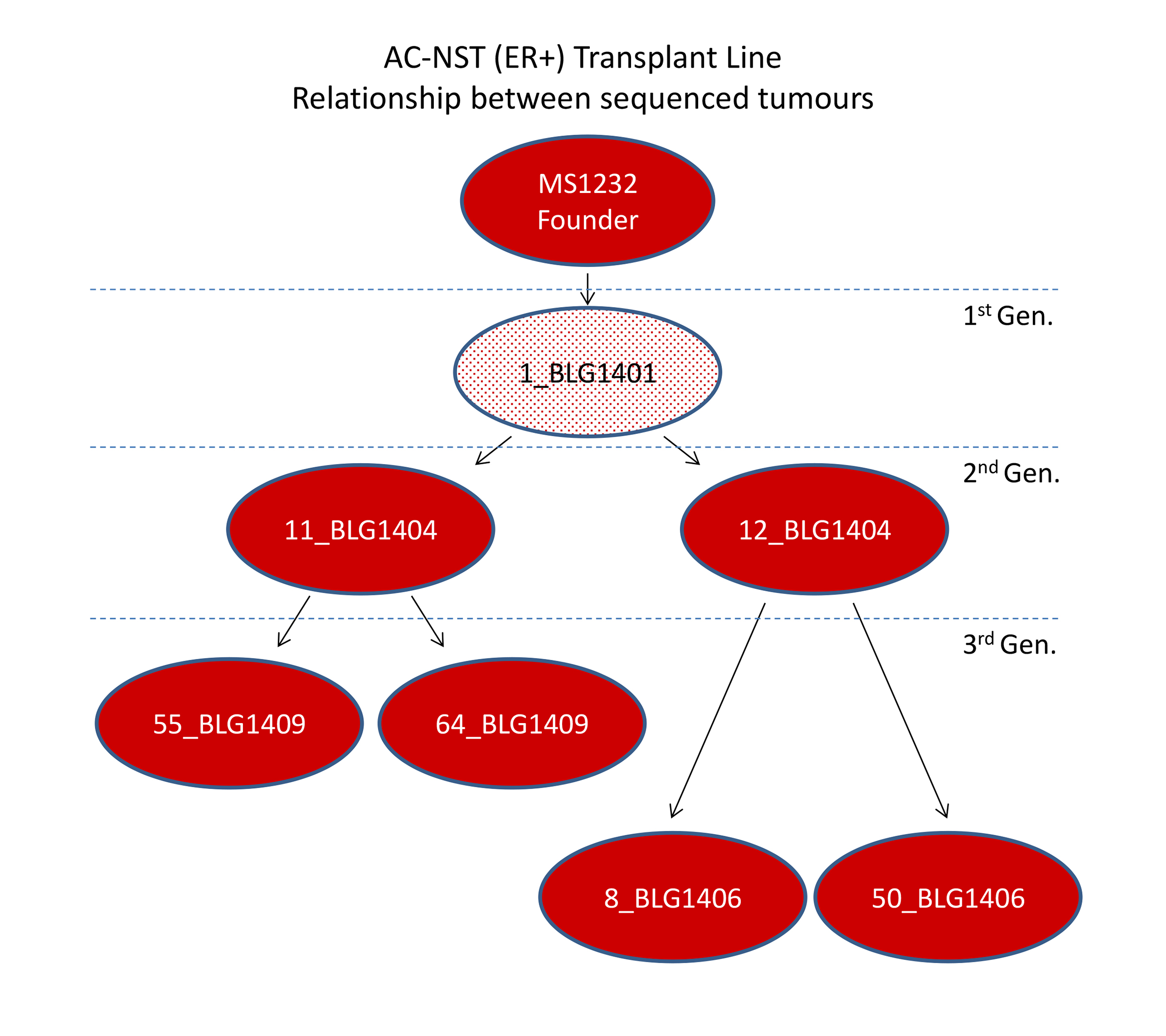

### Supplemental Figure S5

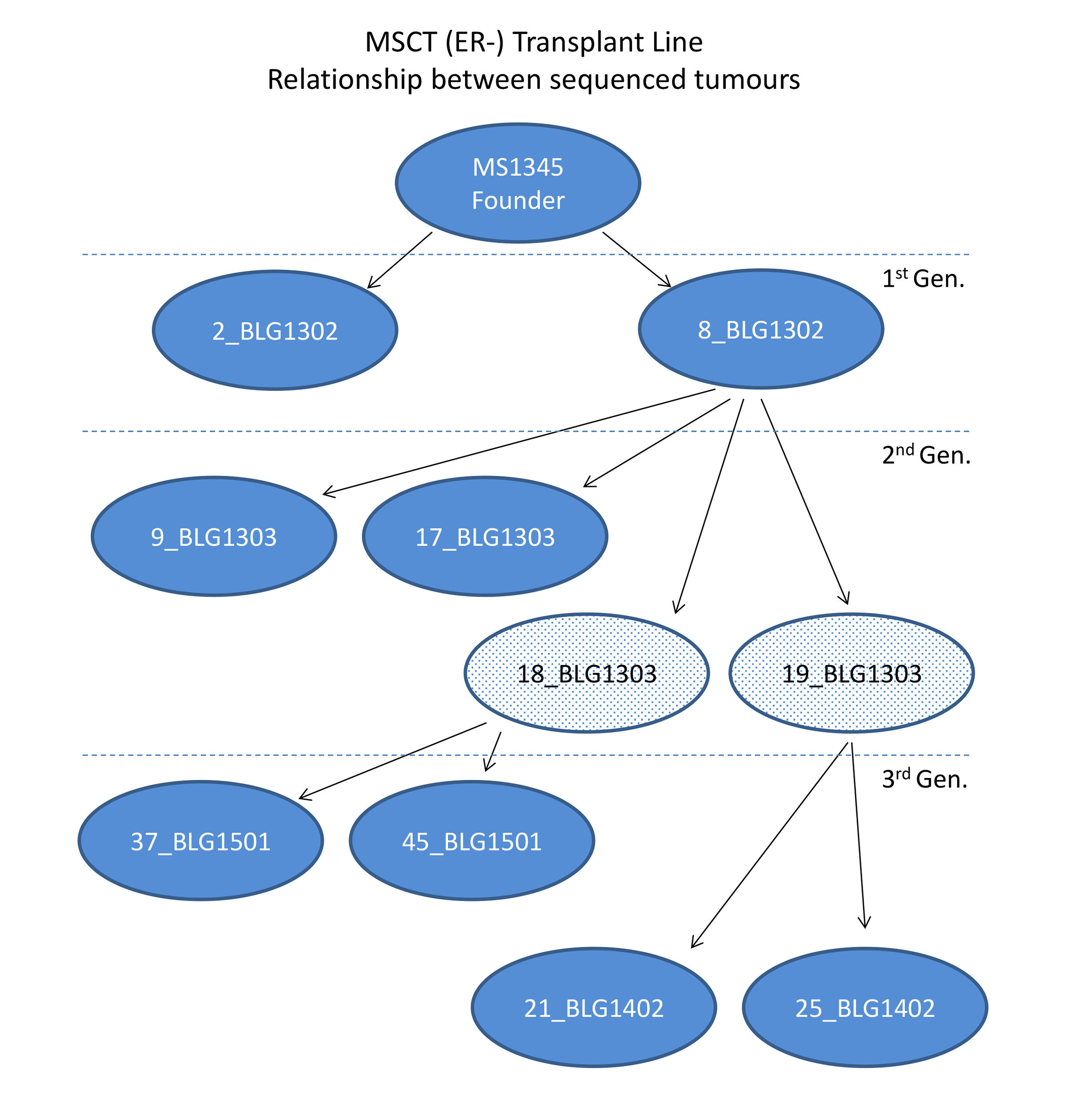
